# Involvement of I-BAR protein IRSp53 in tumor cell growth via extracellular microvesicle secretion

**DOI:** 10.1101/2020.04.20.050492

**Authors:** Hooi Ting Hu, Naoto Sasakura, Daisuke Matsubara, Naoko Furusawa, Masahiro Mukai, Narufumi Kitamura, Takeshi Obayashi, Tamako Nishimura, Kayoko Oono-Yakura, Yosuke Funato, Yasunobu Okamura, Kento Tarao, Yasushi Nakano, Yoshinori Murakami, Kengo Kinoshita, Chiaki Takahashi, Hiroaki Miki, Kohsuke Gonda, Giorgio Scita, Kyoko Hanawa-Suetsugu, Shiro Suetsugu

## Abstract

Cellular protrusions mediated by the membrane-deforming I-BAR domain protein IRSp53 are involved in cell migration, including metastasis. However, the role of IRSp53 in cell proliferation remains unclear. Here, we examined the role of IRSp53 in cell proliferation and found that it acts through secretion. Coculture of gingiva squamous carcinoma Ca9-22 cells and their IRSp53-knockout cells restored proliferation to parental Ca9-22 cell levels, suggesting possible secretion dependent on IRSp53. Notably, the amounts of microvesicle fraction proteins that were secreted into the culture medium were reduced in the IRSp53-knockout cells. The IRSp53-knockout cells exhibited decreased phosphorylation of mitogen-activated protein kinase, suggesting the decrease in the proliferation signals. The phosphorylation was restored by the addition of the microvesicles. In mice xenograft Ca9-22 cells, IRSp53-containing particles were secreted around the xenograft, indicating that IRSp53-dependent secretion occurs *in vivo*. In a tumor mice model, IRSp53 deficiency elongated lifespan. In some human cancers, the higher levels of IRSp53 mRNA expression was found to be correlated with shorter survival years. Therefore, IRSp53 is involved in tumor progression and secretion for cellular proliferation.

## Introduction

The plasma membrane contains various protrusions and invaginations, which are most likely involved in endocytosis or cell migration. These membrane structures are known to be remodeled, at least partly, by Bin/Amphiphysin/Rvs (BAR) domain-containing proteins (Daumke et al., 2014; Doherty and McMahon, 2009; Nishimura et al., 2018; Simunovic et al., 2015; Suetsugu et al., 2014). The BAR domain functions as a scaffold for membrane curvature, in which rigid protein structures mold the membrane (Frost et al., 2008; Frost et al., 2009).

Insulin receptor substrate 53 (IRSp53) and missing-in-metastasis protein share the IRSp53-MIM homology domain (IMD), which is also known as the inverse (I)-BAR domain (Millard et al., 2005; Suetsugu et al., 2006a; Suetsugu et al., 2006b). Therefore, IRSp53 is also known as BAR/IMD domain-containing adaptor protein 2. Alternatively, it is also known as brain-specific angiogenesis inhibitor 1-associated protein 2 (BAIAP2) (Oda et al., 1999).

The N-terminal domain of IRSp53 was first characterized as IMD or Rac binding domain (RCB), which is suggested to bind actin filament and small GTPase Rac (Miki et al., 2000). The binding to small GTPases at this region has also been reported for IRTKS, a paralog of IRSp53 (Millard et al., 2007). IRSp53 and IRTKS are essential genes as their double knockouts exhibit embryonic lethality (Chou et al., 2017). However, structural analysis of the N-terminal region revealed the dimeric helix bundle structure of the IMD, which was the kinked version of the membrane bending BAR domain for adapting the inverted curvature; therefore, the IMD domain is now commonly called the I-BAR domain {Scita, 2008}. The BAR domains of endophilin and amphiphysin contain a concave membrane-binding surface and can thus form a spiral on the surface of membrane tubules corresponding to the plasma membrane invaginations of clathrin-coated pits and others. In contrast, the I-BAR domain has a convex membrane-binding surface, which enables adaptation to the membrane curvature of plasma membrane protrusions, including filopodia (Saarikangas et al., 2009; Sudhaharan et al., 2019; Suetsugu et al., 2006b).

IRSp53 contains several additional domains/motifs, such as the Cdc42- and Rac-interactive binding (CRIB) motif and Src-homology 3 (SH3) domain (Ahmed et al., 2010; Govind et al., 2001; Krugmann et al., 2001; Miki et al., 2000; Scita et al., 2008). IRSp53 was also found to be capable of binding WAVE/SCAR through its SH3 domain (Miki et al., 2000; Oikawa et al., 2013), and the phosphorylated form of IRSp53 is suggested to bind 14-3-3 proteins, resulting in autoinhibition of membrane binding (Cohen et al., 2011; Kast and Dominguez, 2019a, b; Robens et al., 2010).

Filopodia are membrane protrusions that are thought to be mainly regulated by Cdc42. IRSp53 is reportedly involved in cell migration, presumably through filopodia formation (Ahmed et al., 2010; Govind et al., 2001; Krugmann et al., 2001; Scita et al., 2008). However, the role of IRSp53 in cellular proliferation remains unclear. In this study, we generated IRSp53-knockout gingival cancer cells and reduced the amount of IRSp53 by siRNA treatment and found that IRSp53 is involved in the proliferation of the cells. Furthermore, coculture experiments and the tumor implants in mice suggested that the IRSp53-containing particles were secreted from the cells to promote cellular proliferation, suggesting that IRSp53-mediated particle release plays important roles in intercellular communication for cell proliferation. The involvement of IRSp53 in cancer progression has also been suggested based on the lifespan of mice and from cancer genome studies.

## Results

### IRSp53 and proliferation

To delineate the role of IRSp53 in cancer cells, we examined the expression levels of IRSp53 in various cell lines using western blotting (Figure 1A). Among the cells examined, we found that the levels of IRSp53 expression were high in the human gingival squamous carcinoma cell line Ca9-22 (Figure 1A).

**Figure 1.**
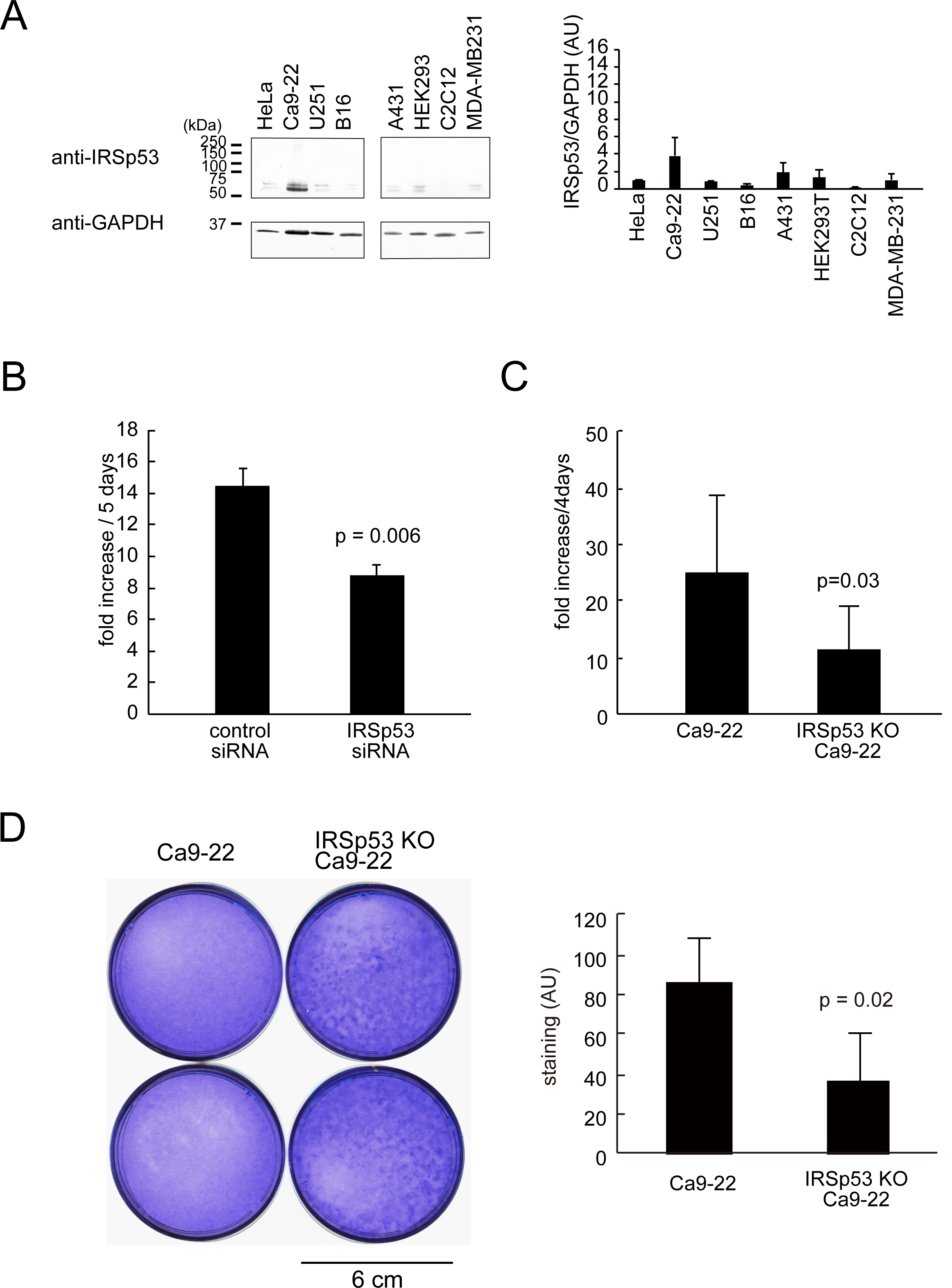
IRSp53 and proliferation. (A) Western blotting of various cell lysates for the amounts of IRSp53 and GAPDH. The quantification of the amount of IRSp53 relative to GAPDH is shown on the right. (B) The fold increase of the cell number of the gingival squamous carcinoma, Ca9-22 cells that are treated with control or siRNA for IRSp53 during 5 days of culture. n=6. (C) The fold increase of the cell number of the gingival squamous carcinoma, parental or IRSp53-knockout Ca9-22 cells during 4 days of culture. n=4. (D) Focus assay of Ca9-22 cells and IRSp53-knockout Ca9-22 cells. Four independent IRSp53-knockout cell lines were analyzed. The cells were seeded on a 6-cm dish, and the cells were stained with crystal violet after 3 weeks. The staining was quantified and has been shown on the right. n=2-5. In (B-D), statistical significance was examined to the Ca9-22 cells using Student’s t-test.

Next, the amount of IRSp53 protein in Ca9-22 cells was reduced by siRNA treatment. Interestingly, the IRSp53-siRNA-treated cells exhibited reduced proliferation compared to that of control-siRNA-treated cells (Figure 1B). Likewise, the CRISPR/Cas9-mediated knockout of IRSp53 also exhibited reduced proliferation compared to that of parental wild-type cells (Figure 1C).

Cell proliferation relates to tumor formation, which is often assessed via focus assays on culture plates. If the cells have tumor forming abilities, the foci of cells will appear in this analysis. Interestingly, parental Ca9-22 cells exhibited foci, but the foci were greatly reduced for the IRSp53-knockout Ca9-22 cell cultures, as examined by crystal violet staining to visualize the entire protein and nucleic acid of the cells (Figure 1D). The staining intensities were significantly different between the parental and knockout cells, confirming that IRSp53 knockout reduced foci formation, presumably via reduction in cell proliferation.

### IRSp53 and possible indirect intercellular communication

IRSp53 is reportedly involved in the formation of cellular filopodia. However, the mechanism via which IRSp53 is involved in cell proliferation remains unclear. To examine whether the role of IRSp53 in cell proliferation is cell autonomous or not, we cocultured the IRSp53-knockout cells with parental Ca9-22 cells. We used the cell culture insert of two wells to seed the cells side by side, with 500μm+/-100μm cell-free gap in between them at the removal of the insert. Without extensive mixing of the cells in the separate two wells, we aimed to examine the possible indirect interaction between the parental cells and IRSp53-knockout cells (Figure 2A). When the two wells of the insert were seeded with the same cells, the knockout cells exhibited reduced occupancy on the plate compared to that of the parental wild-type cells after 7 days of culture (Figure 2B, C), which was consistent with the cell proliferation results (Figure 1C). Remarkably, when the well of knockout cells was sided by the well of parental cells, they exhibited similar growth as that of both wells of parental cells (Figure 2B, C). Therefore, IRSp53 could be involved in intercellular communication between the cells for cellular proliferation.

**Figure 2.**
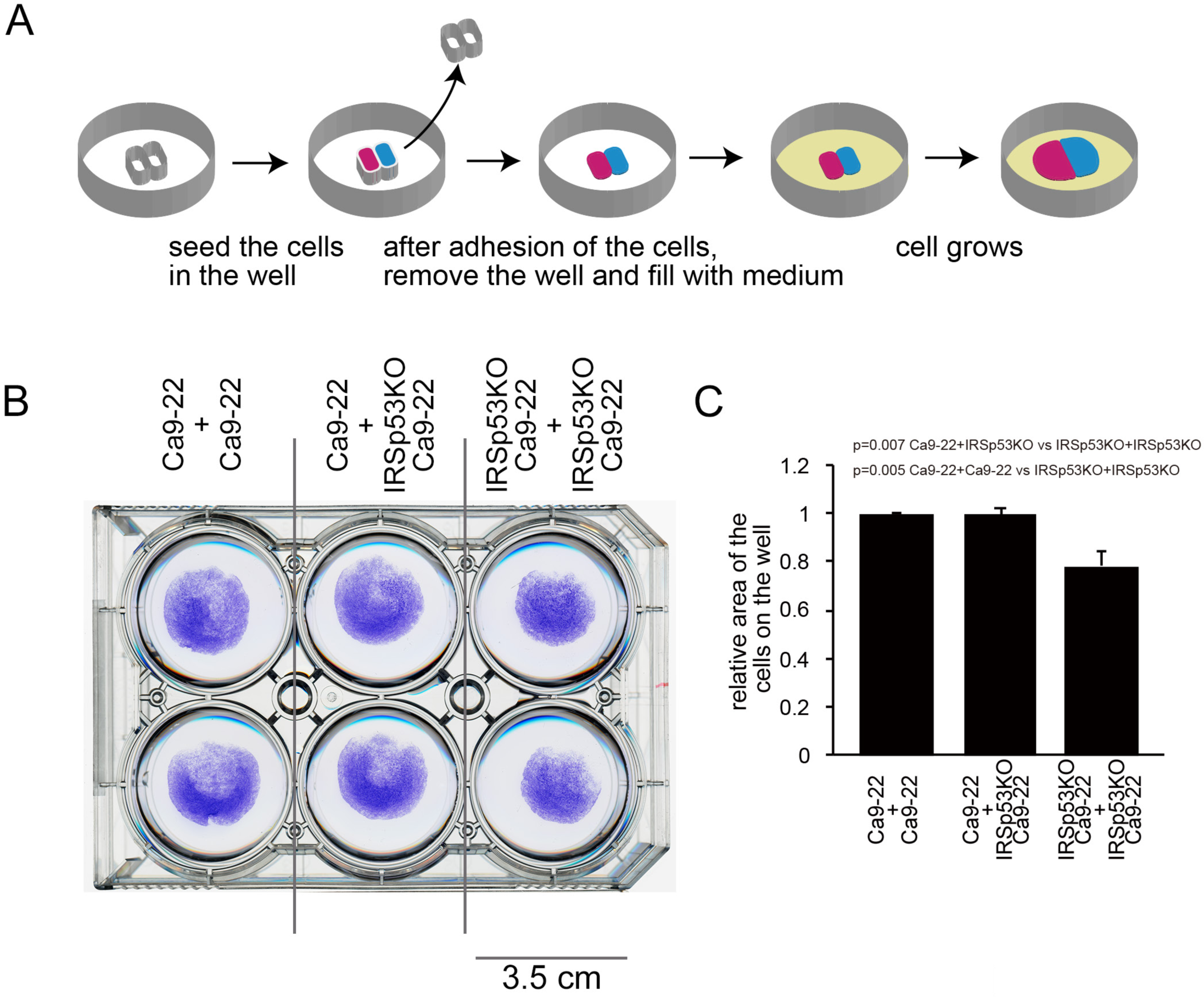
Restoration of proliferation by coculture. (A) Schematic illustration of coculture of the parental and IRSp53-knockout Ca9-22 cells using the cell culture insert. (B) The cells in the plate stained by crystal violet after 7 days of culture. Scale bar: 3.5 cm. (C) The relative area occupied by the cells after 7 days of culture after culture insert removal. Statistical significance was examined using Student’s t-test. n=3.

### Proliferation signal is enhanced by IRSp53-dependent microvesicles

IRSp53 is reportedly involved in filopodia formation. However, the gap between the wells of insert might limit cell-cell direct contact and seems to be too distant for the filopodia to reach the cells at the opposite well. In contrast, secretions from cells, including exosomes and microvesicles, could contribute to cell proliferation (Colombo et al., 2014; Raposo and Stoorvogel, 2013; Tkach and Thery, 2016; van Niel et al., 2018). Exosomes (50–150 nm in diameter) and microvesicles (50–1000 nm in diameter) are membrane-encapsulated vesicles that are thought to originate from the endosome and plasma membrane, respectively (van Niel et al., 2018). Previous studies have reported that microvesicles could facilitate cell–cell communication in cancer cells (Al-Nedawi et al., 2008). To examine the possible roles of microvesicles and exosomes in the promotion of the proliferation of IRSp53-knockout cells, we first obtained the microvesicle and exosome fractions from the culture media of parental Ca9-22 cells and IRSp53-knockout Ca9-22 cells (Figure 3A). IRSp53 was enriched in parental Ca9-22 cell microvesicle and exosome fractions (Figure 3A). Then, we examined the secretion of the cell-surface protein human leukocyte antigen (HLA) (Huet et al., 1980), which is specific to humans and is one of the major classes of major histocompatibility complex (MHC), from the parental Ca9-22 cells and IRSp53-knockout cells. The amount of HLA in the microvesicle fraction was decreased in the IRSp53-knockout cells, whereas that in the exosome fraction was not altered via IRSp53 knockout (Figure 3A, B). Moreover, when IRSp53 was being knocked out, Annexin A1, a marker of microvesicles (Jeppesen et al., 2019), was reduced in the microvesicle fraction, indicating that the loss of IRSp53 could reduce the secretion of microvesicles from the cell (Figure 3A, B). The exosome marker CD81 was not detected in the microvesicle fraction, indicating the non-exosomal characteristics of the vesicles (Figure 3A).

**Figure 3.**
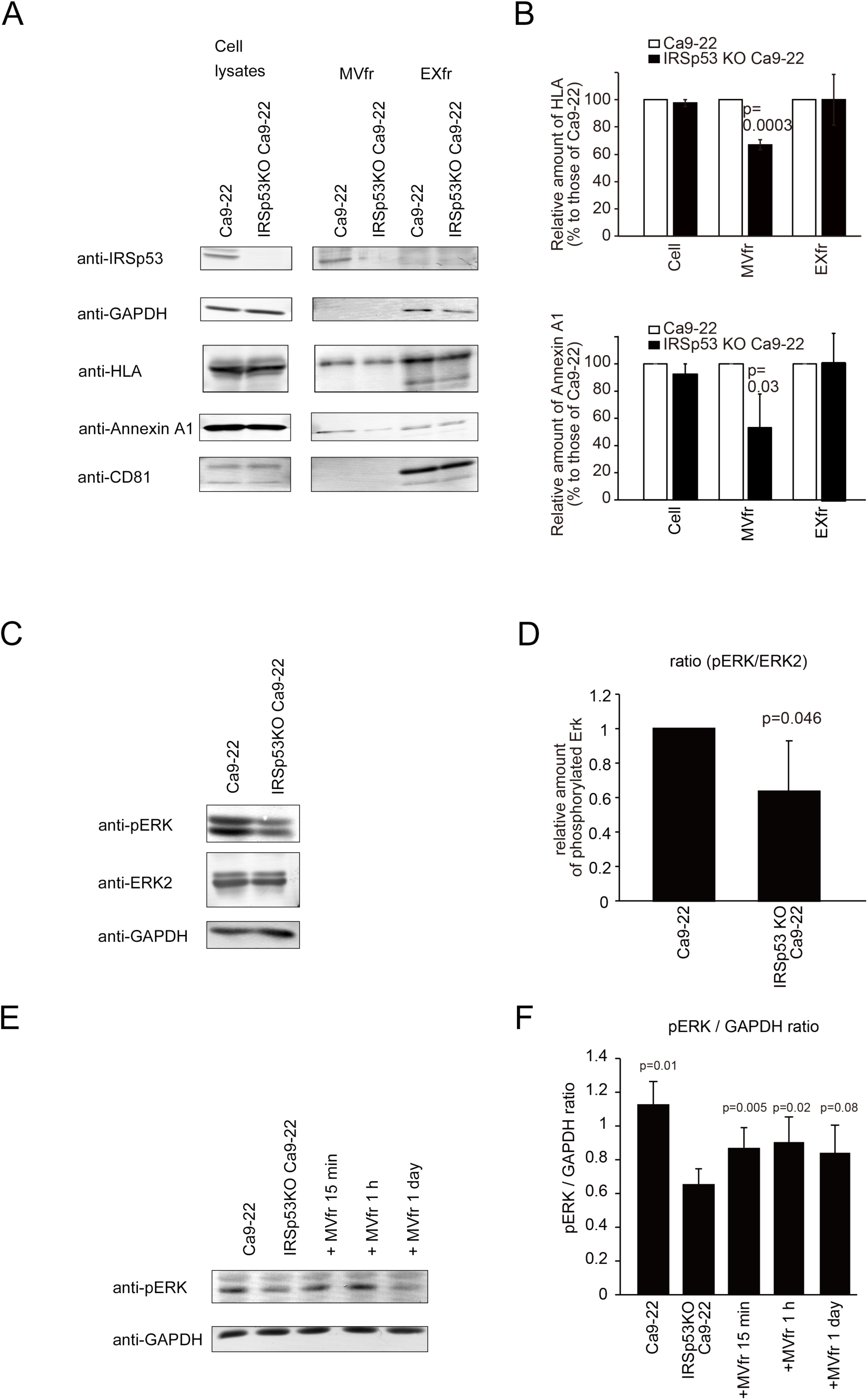
Activation of Erk by microvesicles dependent on IRSp53. (A) Microvesicle fraction (MVfr) and exosome fraction (EXfr) of the parental and IRSp53-knockout Ca9-22 cells. The amounts of IRSp53, GAPDH, HLA, Annexin A1, and CD81 in cell lysates, MVfr, and EXfr from the same numbers of cells were analyzed. (B) Quantification of HLA and Annexin A1 from (A), relative to the parental Ca9-22 cells. Statistical significance was examined to the Ca9-22 cells using Student’s t-test. n=3. (C) Western blot of Erk2 and phosphorylated Erk2 of the parental and IRSp53-knockout Ca9-22 cells. (D) Quantification of (C) relative to the parental Ca9-22 cells. Statistical significance was examined to the Ca9-22 cells using Student’s t-test. n=3. (E) Western blot of Erk2 and phosphorylated Erk2 of the parental and IRSp53-knockout Ca9-22 cells treated with microvesicles from parental Ca9-22 cells. (F) Quantification of (E). Statistical significance was examined to the IRSp53-knockout Ca9-22 cells using Student’s t-test. n=6.

The degree of proliferation signals was then examined based on the extent of phosphorylated mitogen-activated kinase Erk (Stork and Schmitt, 2002). Erk phosphorylation was higher in the parental cells than in the IRSp53-knockout cells (Figure 3C, D). Interestingly, the application of the microvesicle fraction from the parental cells increased the amount of phosphorylated Erk2 of the IRSp53-knockout Ca9-22 cells (Figure 3E, F), suggesting that the microvesicles released from the parental Ca9-22 cells promoted the proliferation of IRSp53-knockout Ca9-22 cells.

### Release of IRSp53 microvesicles from Ca9-22 cells *in vivo*

To obtain insights into the secretion mediated by IRSp53 *in vivo*, we transplanted Ca9-22 cells into immunodeficient mice. Ca9-22 cells subsequently grew into tumors and were fixed. There appeared to be no apparent differences in the growth of the IRSp53-knockout and control Ca9-22 cells in mice, presumably because the surrounding tissue contained IRSp53.

Because IRSp53 was involved in the secretion, we stained the tumor with antibodies specific to human IRSp53, and human HLA was used as a marker for human Ca9-22 cell-derived particles. These human-protein specific antibodies do not bind to the proteins in mice. A secondary antibody coupled with phosphor-integrated dot (PID) (Gonda et al., 2017) was used to enhance the detection of human-cell-derived IRSp53. The IRSp53 antibody stained the parental Ca9-22 cells that were transplanted in mice but did not stain the mice tissues and the IRSp53-knockout cells (Figure 4A, B). The non-binding of the anti-HLA antibody to the proteins in mice was confirmed. Observation under high magnification revealed that particles containing human IRSp53 were found in the vicinity of human Ca9-22 cells, i.e., in the surrounding tissues derived from mice (Figure 4C). Interestingly, these particles sometimes contained human-specific HLA, indicating that these particles were derived from transplanted human cells (Figure 4C, D). The particles were secreted from the cell mass of transplanted Ca9-22 cells, which were absent in IRSp53-knockout cell transplants (Figure 4C, D). Therefore, IRSp53 was strongly suggested to be involved in the formation of particles that were released from the Ca9-22 cells (Figure 4E, F).

**Figure 4.**
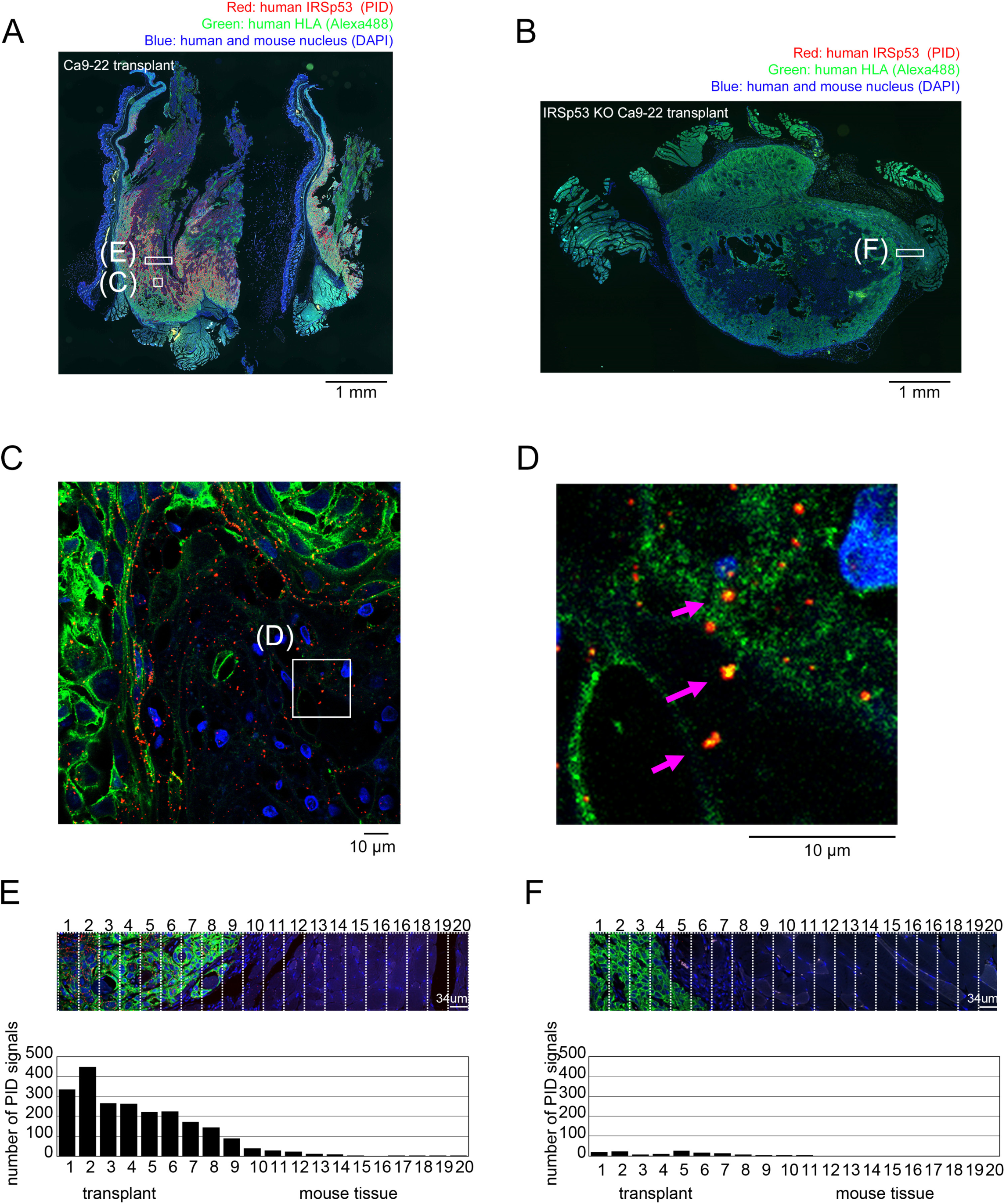
IRSp53 particle release from Ca9-22 cells *in vivo*. (A, B) Transplantation of IRSp53-knockout Ca9-22 cells into immunodeficient NOD/SCID mice that were stained with antibodies specific to human IRSp53 (red) and human HLA (green). The IRSp53 was then further visualized by the bright dye (PID), and the HLA was visualized with Alexa dye. The nuclei were also stained with DAPI (blue), which indicated the nuclei of both human and mouse cells. The images were observed by widefield microscopy. The rectangles indicate the region analyzed in (E, F). The box indicates the region enlarged in (C). (C) The cell-scale view of the transplant of Ca9-22 cells into the mice stained with the antibodies specific for human IRSp53 and HLA. The nuclei were also stained with DAPI. The image was observed by confocal microscopy. (D) The enlargement of the area marked in (C). (E, F) The distribution of HLA containing particles inside (xenograft) and outside (mouse tissue) of the parental Ca9-22 cell (E) or the IRSp53-knockout cell (F) transplant.

### IRSp53 and tumor in mice and humans

Ca9-22 cells contain mutations in the genome-stabilizing protein tp53 (Muller and Vousden, 2013; Sherr and McCormick, 2002), which is often mutated in tumors such as head and neck cancer (Boyle et al., 1993; Somers et al., 1992). To examine the role of IRSp53 in tumor progression, we generated mice deficient in both IRSp53 and tp53. tp53-deficient mice have a shorter lifespan because of the frequent occurrence of tumors (Harvey et al., 1993; Merritt et al., 1994). The heterozygous deficiency of IRSp53 was resulted in the reduction in the amount of IRSp53 proteins as examined by the western blotting of thymus and brain tissue lysates (Figure 5A). Interestingly, the heterozygous deficiency of IRSp53 was sufficient to prolong the lifespan in female mice, suggesting that the amount of IRSp53 influences the lifespan of mice (Figure 5B). Such a difference was gender specific and detected only in female mice. The number of female mice that were deficient in both IRSp53 and tp53 were limited due to the known embryonic lethality of IRSp53 mice {Chou, 2017 #67}. Hence, we exclude them for the lifespan analysis. Apparently, there were no differences in the types of tumor or in their occurrence comparing tp53^-/-^ mice and compound IRSp53^-/+^/tp53^-/-^ mice, suggesting that lifespan increase was due to a general reduced tumor burden by the reduced interactions between the cells and the tissues.

**Figure 5.**
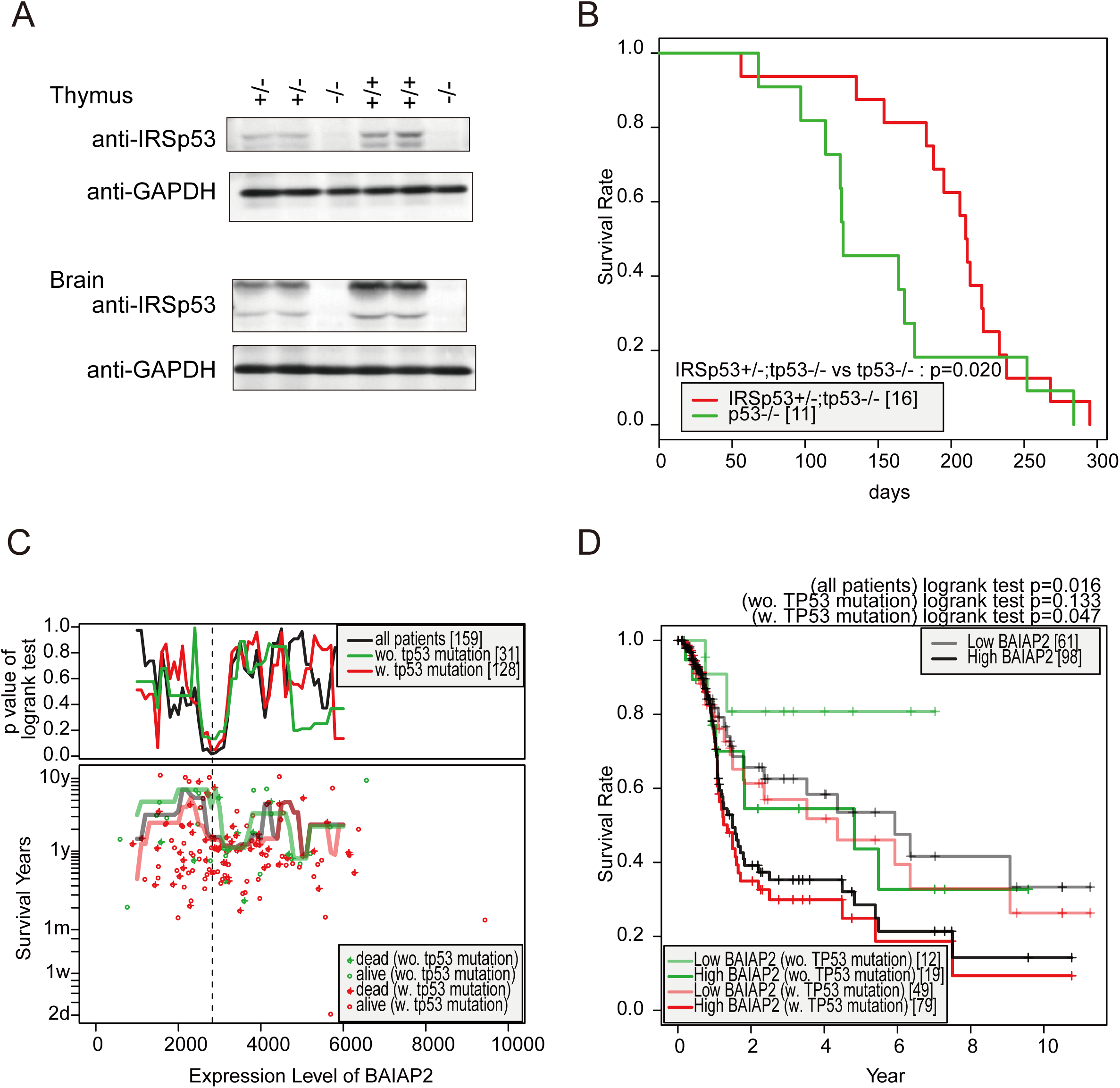
IRSp53 and tumor in mice and human. (A) The amount of IRSp53 in mice. The littermate of the crossing of IRSp53 heterozygous mice were analyzed at P9 for their amount of IRSp53 and GAPDH at brain and thymus by western blotting. (B) Kaplan–Meier analysis of heterozygous IRSp53 and homozygous tp53-knockout female mice. The number of mice in each group is shown in brackets. The p value by log-rank test is shown. (C) The lifespans of the patients having head and neck cancer depending on the amounts of IRSp53 mRNA using TCGA. The mRNA amounts (RNA Seq V2 RSEM) from 1000 to 6000 with 100 intervals for each of the all 159 patients, for the 31 patients without TP53 mutation, and for the 128 patients with TP53 mutation were used for analysis (upper panel). For visual interpretation, the median survival time of the patients having a particular range (1000 in width) of IRSp53 mRNA amount was calculated with every 100 intervals from 1000 to 6000 (lower panel). In the case that the median survival time was undecidable due to the censored data in longer survival time, it was conservatively approximated by regarding the last censored event as death event. (D) Kaplan–Meier analysis of the head and neck cancer patients. Kaplan-Meier plot with the threshold of IRSp53 expression level, 2800, was performed between patient with high expression IRSp53 (deep colors) and that with low expression (pale colors) for each of the all patients (black), the patients without TP53 mutation (green), and patients with TP53 mutation (red). The number of patients in each group is shown in brackets. The p value by log-rank test has been shown.

We then examined the relation of the amount of IRSp53 expression levels and the lifespans in human cancers using The Cancer Genome Atlas (TCGA). The lifespans of patients with high and low IRSp53 expression were examined using various levels of IRSp53 mRNA as the threshold value between the two populations. Among the cancers listed in TCGA, patients with head and neck cancer, in which gingival carcinoma is included, had significantly longer lifespans when their amounts of IRSp53 expression were low (Figure 5C). The moving median lines of the survival time displayed the general trend that patients having lower amounts of IRSp53 mRNA showed longer lifespans (Figure 5C, lower panel). The dependency on the threshold levels of IRSp53 was similar among the three groups tested (all 159 patients, 31 patients without TP53 mutation, and for the 128 patients with TP53 mutation) (Figure 5C, upper panel). Kaplan–Meier plots with the optimal threshold of the mRNA amount (2800) are shown in Figure 5D. Patients with low amounts of IRSp53 mRNA (pale colors) had significantly longer lifespans than those with high amounts of IRSp53 mRNA (darker colors) for each of the all patients (black), as well as for the patients without (green) or with (red) tp53 mutation. Thus, the high levels of IRSp53 impact on lifespan regardless of tp53 status, reinforcing the concept that IRSp53 is an important determinant of survival in patient with head and neck cancer. The differences between males and females could not be tested because relevant data were not available. The involvement of the amounts of IRSp53 in the lifespan of cancer patients was found for several other cancers in TCGA, which include glioblastoma with tp53 mutation (Figure S1). Altogether, these data suggested that elevated levels of IRSp53 expression could promote tumor development or progression both in mice and humans.

## Discussion

We found that IRSp53 could be involved in the proliferation of cells via possible secretion of growth enhancing factors. Importantly, the secretion of IRSp53-containing particles was observed in cancer cell xenografts, suggesting that IRSp53-dependent secretion occurs in physiological situations.

These particles appeared to be IRSp53-dependent microvesicles, which might be capable of long-distance communication in cell proliferation. Microvesicles can promote cell proliferation (Gupta et al., 2017; Gurunathan et al., 2019; Muralidharan-Chari et al., 2010). In this study, the mechanisms underlying the enhancement of proliferation by IRSp53-mediated vesicles have not been clarified. Further studies are warranted to characterize the proteins and molecules in IRSp53-containing vesicles.

Microvesicles could be generated via direct scission of the plasma membrane (Colombo et al., 2014; D’Souza-Schorey and Clancy, 2012; Jeppesen et al., 2019; Kowal et al., 2016; van Niel et al., 2018). However, no molecular mechanism for this has been suggested. Because the BAR domain protein endophilin was reported to promote intracellular vesicle formation (Gallop et al., 2006; Simunovic et al., 2017), it might be reasonable to hypothesize that the I-BAR domain of IRSp53 can promote vesicle formation as well. IRSp53 is reportedly involved in the formation of filopodia, which are plasma membrane protrusions that inversely direct membrane invagination. Indeed, such plasma membrane scission resembles virus budding, and IRSp53 has been suggested to be involved in the release of human immunodeficiency virus and pseudorabies virus from the plasma membrane (Thomas et al., 2015; Yu et al., 2019). Thus, one candidate mechanism for microvesicle generation is direct scission of the filopodia, in a manner analogous to that of endophilin-mediated promotion of scission of the invaginations in cooperation with other factors, including physical forces (Simunovic et al., 2017), because there are supposedly abundant physical forces present on the outside of cells.

Tumors are known to frequently occur in tp53-deficient mice (Jeppesen et al., 2019). Interestingly, I-BAR domain proteins might be involved in tumor development when the amount of tp53 is reduced. The relationship between I-BAR domain proteins and tumors has been reported in IRTKS-knockout mice (Huang et al., 2018). IRTKS is a paralog of IRSp53, and the two share similar domain structures. In case of IRTKS, tp53-heterozygous mice deficient in IRTKS had longer lifespan than the tp53-heterozygous mice having IRTKS (Huang et al., 2018).

Aggressive cancer cells have been demonstrated to be more persistent and have more filopodia-like protrusions as compared to nonaggressive cancer cells (Jacquemet et al., 2015; Shibue et al., 2013). In this study, although no clear differences were identified in the types of tumors as well as the occurrence of metastasis in the mice, the presence of IRSp53, which could initiate filopodia formation, was possibly related to cancer cell aggressiveness and poor prognosis. Furthermore, the phenotype specific to female mice might suggest something related to the secretion, as secretion might differ between female and male mice. If IRSp53 is involved in secretion, it may possibly affect the interaction of the tumor with the surrounding organs. Cancer cells can interact with their surrounding organs via secretion and by creating a tumor-promoting microenvironment (Egeblad et al., 2010). Therefore, IRSp53-mediated secretion might be related to the degree of tumor burden. The high metabolic tumor burden, such as increase in tumor volume, was associated with poor prognosis and low survival (Kelemen et al., 2012; Zhang et al., 2014).

Secretion may vary based on cell types and organs in which I-BAR proteins are expressed; therefore, IRSp53-dependent secretion might not be found in all types of cells or organs. Further studies are required for clarifying the mechanisms underlying IRSp53-dependent secretion and its relevance in various types of cells and organs as well as in tumorigenesis. However, our study demonstrated that the involvement of IRSp53 in microvesicle release could occur in physiological situations.

## Methods

### Cell culture, siRNA treatment, and CRISPR/Cas9-mediated genome editing

The cells were cultured in DMEM supplemented with 10% FBS. Transfection of Ca9-22 cells (2.5 × 10^5^ cells/well of a 6-well plate) with siRNA (6 μl, 20 μM) was performed using Lipofectamine RNAi Max (1 μl) (IRSp53 siRNA: 5’-AGUACUCGGACAAGGAGCUGCAGUA-3’; control siRNA: 5’-AGUGGCUAACAGAGGCGUCACAGUA-3’.)

The CRISPR/Cas9 system was used, as described previously (Mashiko et al., 2013). The guide RNA targeting the first exon of IRSp53 (CCATGGCGATGAAGTTCCGG) was designed using the server http://crispr.mit.edu (Hsu et al., 2013) and inserted into the pX330 vector (Mashiko et al., 2013). After transfection, the cells were cloned by monitoring the GFP fluorescence from the reporter plasmid pCAG-EGxxFP with the IRSp53 genome fragment using a fluorescence activated cell sorter [FACSAria (BD)] (Hanawa-Suetsugu et al., 2019).

### Focus Assay

Ca9-22 cells and IRSp53 knockout Ca9-22 cells were cultured in DMEM supplemented with 10% calf serum (Invitrogen). Cells (1 × 10^5^) were seeded into a 6-cm dish. The cells were then cultured for 3 weeks. The culture medium was replaced every 2–3 days. Focus formation was analyzed using crystal violet staining.

### Coculture experiments

Ca9-22 cells and IRSp53 knockout Ca9-22 cells were cultured in DMEM supplemented with 10% calf serum (Invitrogen). Cells (5 × 10^5^) were seeded into a well of culture-insert 2 well (ibidi, 80209). The cells were then cultured for 7 days without replacement of the medium and visualized using crystal violet staining.

### Microvesicle and exosome preparation

The culture medium atop cells was collected and supplemented with 0.1 mM PMSF and 5 mM EDTA. Cellular debris was removed by centrifugation at 3,000 *×g* for 10 min at 4°C. The medium was then centrifuged at 16,500 *×g* for 20 min at 4°C to pellet the microvesicles. The supernatant was then collected and subjected to exosome preparation. The pellet of microvesicles was washed once with ice-cold PBS. The resulting microvesicles were suspended in PBS supplemented with β-mercaptoethanol for analysis.

For the preparation of exosomes, the supernatant collected as mentioned above and centrifuged at 120,000 *×g* for 70 min at 4°C. The pellet was washed once with ice-cold PBS. The resulting exosomes were suspended in PBS supplemented with β-mercaptoethanol for analysis.

### Western blotting

Western blotting was performed using anti-IRSp53/BAIAP2 (Sigma, HPA023310), anti-GAPDH (Santa Cruz Biotech, sc-166574), anti-p-ERK(E-4) (Santa Cruz Biotech, sc-7383), anti-ERK2 (D-2) (Santa Cruz Biotech, sc-1647), anti-HLA class I (HLA-A,B,C) (EMR8-5) (MBL, D367-3), anti-Annexin I (EH17a) (Santa Cruz Biotech, sc-12740) and anti-CD81 (B-11) (Santa Cruz Biotech, sc-166029) were used as the primary antibodies. Alkali-phosphatase (AP)-conjugated anti-mouse or anti-rabbit IgG (Promega) as well as AP substrate consisting of 5-bromo-3-chloro-indolyl phosphate (Roche Diagnostics) and 4-nitro blue tetrazolium chloride (Roche Diagnostics) were used for detection. The signals were quantified using ImageJ software (NIH, USA).

### Transplantation of Ca9-22 cells into mice

Ca9-22 and IRSp53 knockout Ca9-22 cells were cultured in DMEM supplemented with 10% calf serum (Invitrogen). Cells (2 × 10^6^ cells/100 μl) were mixed with Matrigel (100 μl) (BD) on ice and hypodermically injected into NOD-SCID mice using a 22-gauge needle. After 4 months, the xenografts were excised, fixed in 3.7% formaldehyde in PBS, and then embedded in paraffin.

### Tissue staining

All paraffin sections were deparaffinized in xylene and hydrated in a series of graded alcohol and distilled water. Antigen retrieval was performed by autoclaving the samples in 10 mM citrate buffer (pH 6.0) for 5 min at 121°C. Then, the samples were immunostained with the anti-IRSp53/BAIAP2 antibody 1D9 (mouse monoclonal, 1:200, OriGene) and anti-HLA A antibody EP1395Y (rabbit monoclonal, 1:100, Abcam) at 4°C overnight. The samples were washed with PBS and then incubated with biotinylated anti-mouse IgG secondary antibody (4 µg/ml, abcam) and Alexa Fluor 488 conjugated anti-rabbit IgG secondary antibody (1:250; Thermo Fisher) at 25°C for 30 minutes. After incubation with secondary antibodies, the sample was treated with 0.02 nM PID at 25°C for 2 h. The sample was washed, stained with DAPI (1:10000) for 10 minutes, and mounted in mounting medium (marinol 550cps, Muto pure chemicals).

Fluorescence signals were observed using the widefield fluorescence microscope Keyence BZ-X710 for low magnification images and the confocal microscope Zeiss LSM880 with airyscan for high magnification images. The excitation lengths of the confocal microscope for IRSp53 (PID), HLA (Alexa488), and DAPI were 561, 488, and 405 nm, respectively.

### Mice

The IRSp53-knockout mice were generated as described previously (Sawallisch et al., 2009). The tp53-knockout mice were also generated as described previously (Tsukada et al., 1993). Animal maintenance and animal experiments were performed in accordance with the guidelines of the Committee on Animal Research at Nara Institute of Science and Technology.

### TCGA analysis

A survival analysis on human head and neck squamous cell carcinoma was performed using data provided by TCGA (https://www.cbioportal.org/study/summary?cancer_study_id=hnsc_tcga_pub) (Cerami et al., 2012; Gao et al., 2013), particularly focusing on the effect of the mRNA level of IRSp53 with the presence and absence of the tp53 mutation. Because the amount of mRNA is a continuous measurement, the effect of the threshold to separate higher and lower amounts of IRSp53 mRNA was first investigated as similarly provided in The Human Protein Atlas (Uhlen et al., 2015). Survival analyses were performed using the survival package (Therneau, 2015) in R (version 3.6.2).

## Acknowledgements

We thank Prof. Masahito Ikawa (Osaka University) and Prof. Taro Kawai (Nara Institute of Science and Technology) for the CRISPR/Cas9 system, Ms Ayumu Hashida, Dr Andrea Disanza (IFOM), and all the members of the laboratories for technical assistance and helpful discussions. This work was supported by grants from the Funding Program for Next Generation World-Leading Researchers (NEXT program LS031), JSPS (KAKENHI 26291037, JP15H0164, JP15H05902, JP17H03674, JP17H06006, JP20H03252), JST CREST (JPMJCR1863), Osaka Cancer Research Foundation, the Naito Foundation, the Sumitomo Foundation, the Mitsubishi Foundation, the Sagawa Foundation for Promotion of Cancer Research, and the NAIST Interdisciplinary Frontier Research Project to S.S., JSPS (KAKENHI JP16K07351) to K.H.-S., the Associazione Italiana per la Ricerca sul Cancro (AIRC-IG#18621, and 5XMille #22759), The Ministry of Research and University (PRIN: Progetti di Ricerca di Rilevante Interese Nazionale – Bando 2017#2017HWTP2K) to GS. This work was partly performed in the Cooperative Research Project Program of the Institute of Medical Science, the University of Tokyo, the Research Institute for Microbial Diseases, Osaka University, and the Cancer Research Institute, Kanazawa University.

## Author Contributions

H.T.H., N.S., T.N., K.O.-Y., K.T., K.H.-S. and S.S. performed cell biological experiments. D.M., N.F., M.M., N.K., Y.F., K.O.-Y, K.H.-S., and S.S. analyzed mice and transplanted cells. T.O., and Y.O. analyzed the TCGA data. C.T. and G.S. provided mice. Y.N., Y.M., K.K., C.T., H.M., K.G., G.S., and S.S designed the research and supervised the project. H.T.H, N.F., T.O., and S.S. wrote the manuscript with input from all other authors.

## Author Information

Correspondence and requests for materials should be addressed to S.S.

## Figure Legends

**Supplementary Figure 1.**
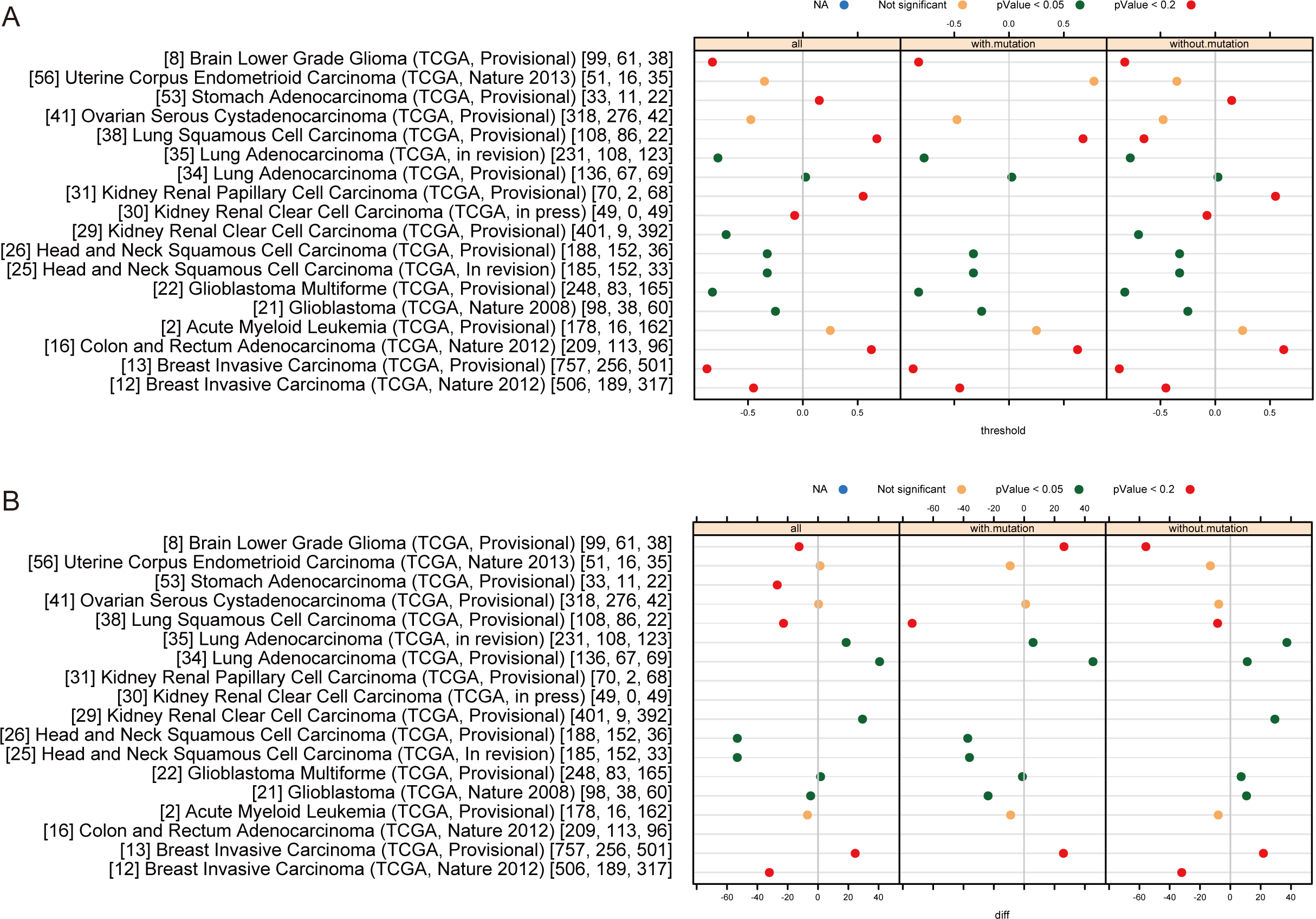
Effect of the amount of IRSp53 mRNA on various types of cancers. To survey potential associations between cancer type and IRSp53 expression, survival analysis was conducted on various types of cancers, the raw data of which were downloaded from TCGA on Jan 14, 2014. The results of the log-rank test between patients with high and low expression of IRSp53 mRNA are shown by (A) optimal threshold of z-scored mRNA level to separate high and low IRSp53 expression and (B) difference of median survival time. Each graph is composed of three panels: results based on all patients regardless of the tp53 mutation status (left), patients with tp53 mutation (center), and patients without tp53 mutation (right). The number of patients for these three categories are shown in the data names with square brackets.

